# Neural correlations are attenuated in the transformation from the frontal cortex to movement

**DOI:** 10.1101/2021.04.29.441941

**Authors:** Noga Larry, Mati Joshua

## Abstract

Correlated activity between neurons can cause variability in behavior across trials. The extent to which correlated activity affects behavior depends on the properties of its translation into movement. We developed a novel method that estimates the contribution of correlations in the frontal eye field (FEF) to pursuit eye movements. We defined a distance metric between the behavior on different trials. Based on this metric, we applied a sequence of shuffles to the neuronal responses, allowing trials to be matched with increasingly distant trials. Correlations between neurons were strongly attenuated when applying even the most constrained shuffle. Thus, only a small fraction of FEF correlations affects the behavior. We used simulations to validate our approach and demonstrate its generalizability over different models. We show that the attenuation of correlated activity through the motor pathway could stem from the interplay between the structure of the correlations and the decoder of FEF activity.

## Introduction

The effect of correlated noise on neural computation has been studied both experimentally and theoretically (Abbott and Dayan, 1999; Khanna et al., 2019; Medina and Lisberger, 2007; Moreno-Bote et al., 2014; Schneidman et al., 2006; Shamir and Sompolinsky, 2004; Zohary et al., 1994). The main focus of these studies has been to identify different conditions in which correlations are decremental or incremental to the amount of information a population carries on sensory stimuli. This framework provides upper bounds on the available information within a network but does not specify whether or how correlated noise propagates to downstream computations. Here we use the eye movement system to study the propagation of correlated noise from a motor area and onwards to behavior.

The activity of single neurons in the motor system is correlated with fluctuations in behavior across trials (Britten et al., 1996; Churchland et al., 2006; Joshua and Lisberger, 2014; Medina and Lisberger, 2007). The independent variability of a single neuron is too small to drive these fluctuations, and therefore the correlation with behavior must arise from the coordinated activity of many neurons (Ben-Shaul et al., 2001; Brody, 1999; Haefner et al., 2013; Musall et al., 2019; Stringer et al., 2019). Figure 1A illustrates an example of how the common coding of variable movements can explain correlations between neurons. In this simplistic model, the activity of each neuron in the population consists of a common variable that is shared with the rest of the population (blue part) and an independent variable (red part). The population activity is read out to produce movement (e.g., the speed of eye movements in a given direction) in a manner that averages out the independent components. Thus, the behavior produced by this simple network is completely determined by the shared component. In this case, the correlation between neurons can be fully explained by the common coding of the behavior. However, in more complex networks, neurons may be correlated beyond the contribution of movement.

**Figure 1:**
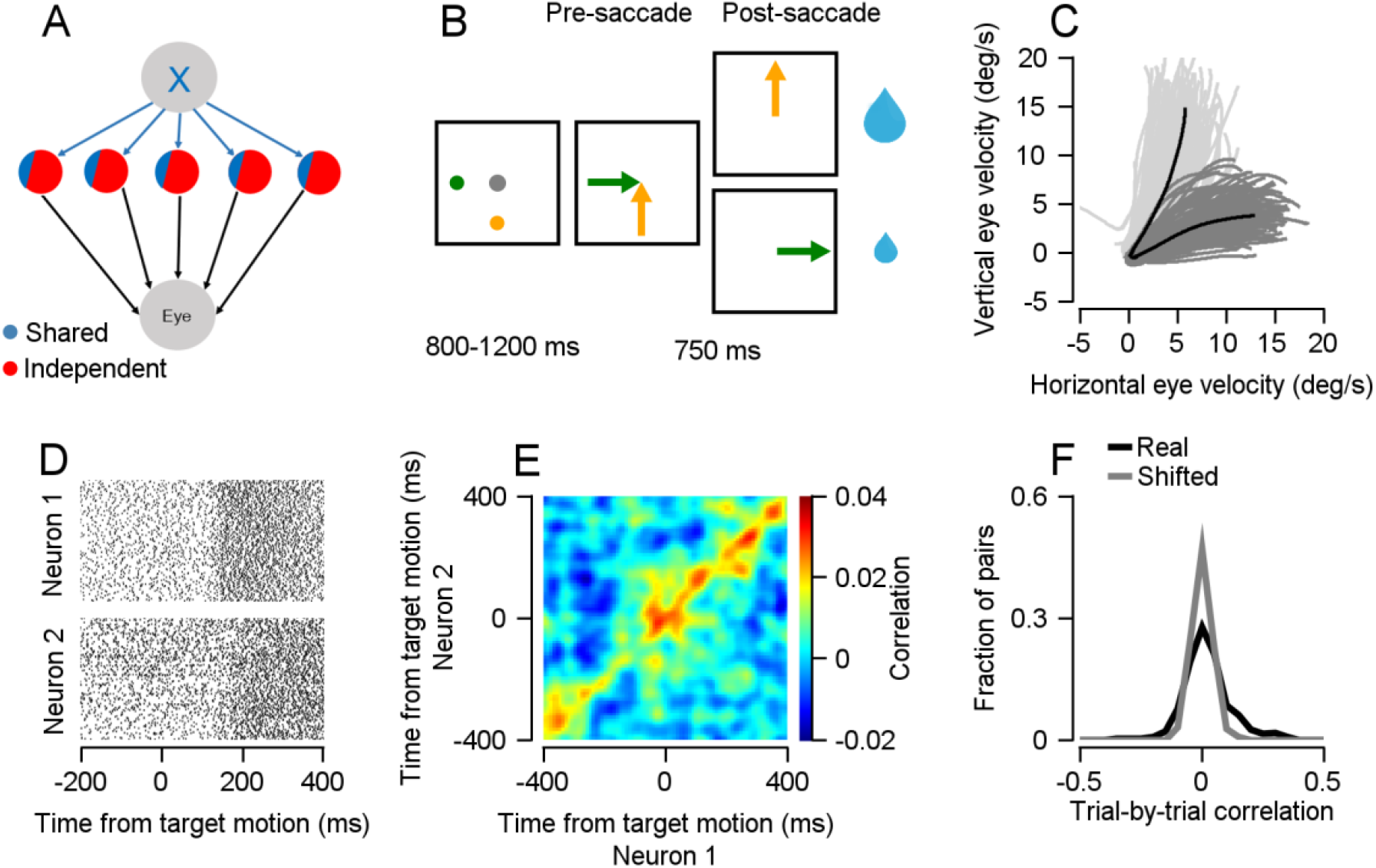
Correlations between neurons recorded simultaneously in a pursuit eye movement task. **A**, A model of a neural population in which correlated variance between neurons is caused by common coding of a movement variable (blue part) and uncorrelated variance by independent noise (red). **B**, Pursuit eye movement task in which the monkey chooses between two targets associated with different reward sizes. **C**, Pursuit eye velocities of individual trials in the 220 ms after target motion onset for trials in which the larger reward target moved along the vertical axis (light grey) or along the horizontal axis (dark grey). The average traces are in black. **D**, Example raster plots of two neurons recorded simultaneously. **E**, The population correlation matrix calculated separately for each condition and averaged over pairs of neurons. The color of each point in the matrix represents the correlation between the response of neuron 1 (horizontal) and neuron 2 (vertical) in time widows aligned to the target motion. **F**, Histograms of the trial-by-trial correlations of pairs of neurons in the 50 to 150 ms after target motion onset (black) and of the correlation calculated after shifting the trials in neuron 1 one trial forward (gray; two-sample Kolmogorov-Smirnov: p=10^-9, n=470).

We examined to what extent correlations between neurons are explained by fluctuations in movement. We developed a novel method to separate neural correlations into correlations that propagate to specific behavior and contribute to its variance, termed *behavior-related correlations* and correlations that do not. We achieved this separation by defining a sequence of shuffles on the neural responses, with decreasing constraints on the similarity of the behavior between shuffled trials. We used the shuffling method to analyze neural correlations in the frontal eye field (FEF) during a smooth pursuit task. Most of the correlations between neurons before and during movement were not expressed in the behavior.

Our approach does not assume a specific model of the relations between neurons and behavior. An imprecise model could result in either an over- or underestimation of behavior-related correlations. By using shuffling, we make only minor assumptions about the coding of behavior by the neurons, making our method generalizable to different models of the FEF. We used simulations to show that our method accurately reflects behavior-related correlation under these models. We then assumed a plausible decoder and demonstrated that correlations that are not expressed in the behavior arise from the interplay of the correlation structure and the decoder. Thus, we used models to describe the conditions in which correlated activity is attenuated as it propagates downstream through the motor system.

## Results

### Neurons in the FEF are correlated during eye pursuit movement initiation

We recorded FEF neurons while monkeys performed an eye movement task in which they were required to track one of two targets based on their reward values (Figure 1B; Joshua and Lisberger, 2012; Raghavan and Joshua, 2017). Each trial began with a fixation period, which was followed by the appearance of two peripheral colored targets. The color of one of the targets was associated with a large reward (e.g., yellow in Figure 1B) and the color of the other target was associated with a small reward (green). The large and small reward targets always appeared on the same axes in each session (for example up and right in Figure 1B). However, we interleaved trials in which their locations were switched. The monkey was required to maintain fixation until the peripheral targets began to move toward the center of the screen at a constant speed. The monkeys typically initiated a smooth pursuit eye movement that was biased toward the target that was associated with the larger reward (Figure 1C). This bias indicated that monkeys used the information about target location and reward to prepare pursuit towards the large reward target (Joshua and Lisberger, 2012). We therefore refer to the presentation of the peripheral targets as the cue. Later in the trial, the monkeys made a corrective saccade which in 99% of the trials brought the position of the eyes close to the large reward target. The monkeys received a reward based on the reward value associated with the target to which they saccaded.

The neurons in our sample responded to the presentation of the peripheral targets as well as to the movement initiation (Raghavan and Joshua, 2017). At both the behavioral and the neuronal level, there was a typical response to target motion that was visible when averaging over repetitions of the same trial. However, at the individual trial level, there was variability both in the behavior (gray traces in Figure 1C) and in the neuronal activity (Figure 1D, differences between trials in the raster plots). The task structure induced variable behavior since the bias of the eye movement toward the larger-reward target differed between trials. We took advantage of this variable behavior to study the trial-by-trial correlations of neurons and their connection to behavior.

Our sample contained 224 pairs of neurons that we recorded simultaneously in two experimental conditions defined by the respective locations of the large and small reward targets. To quantify the trial-by-trial co-fluctuation of these pairs of simultaneously recorded neurons, we calculated the Pearson correlation matrix separately for each condition. Figure 1E shows the population correlation matrix, averaged over conditions, and aligned to the target motion onset. Each point in the correlation matrix represents the Pearson correlation between the response of one neuron (horizontal axis) and the response of the second neuron (vertical axis) in 50 ms bins. Points on the main diagonal show the correlations of simultaneous activity, while off-diagonal points show the correlations of activity that occurs at different times in the trial. On average, neurons were positively correlated along the diagonal of the matrix.

To examine the pairs separately, we calculated the distribution of the average main diagonal correlations following motion onset (Figure 1F, black curve). The distribution had a positive tail, indicating that although the stimuli did not change across trials, the responses of these neurons fluctuated together. To verify that these correlations were not an artifact of slowly occurring changes in firing rates, such as drifts or recording instability, we shifted the order of the trials in one of the two neurons by one trial and recalculated the correlations. The distribution became narrower and symmetric (Figure 1F, grey curve), indicating that the correlations we observed are due to short-term covariation between trials.

### Matching behaviorally similar trials to estimate the contribution of trial-by-trial correlations to behavior

Variability that is shared across a population can influence behavior, causing it to differ on a trial-by-trial basis (Figure 1A). It is, however, possible that part of the correlated components in a population’s activity does not propagate to the behavior due to the properties of the translation of its activity into movement. Next, we aimed to assess the fraction of trial-by-trial correlations between neurons in the FEF related to trial-by-trial variation in pursuit behavior.

Our analysis is based on the following logic: If all the neural correlations are a result of a shared component that reaches behavior (e.g., Figure 1A), we could hypothetically take one of the neurons in a pair and replace its firing rate on individual trials with its firing rate on trials with identical behavior. As the only component of the first neuron’s variance that is shared with the second neuron is equal on behaviorally identical trials, this shuffle should not affect the correlation between the neurons. Of course, it is impossible to find two trials with exactly the same behavior. However, at least part of the correlation should remain when matching behaviorally similar trials. Correlations that do not affect behavior would not be preserved following the shuffle. Taking this logic a step farther, if the shared component reaches behavior, the correlations should decrease as the matched trials become more distant.

To estimate the fraction of correlations that can be explained by common coding of the behavior (*behavior-related correlations*), we applied a sequence of shuffles to each pair of simultaneously recorded neurons in each experimental condition. In each shuffle in the sequence, we allowed the trials to be shuffled with trials that were less similar behaviorally. To quantify the similarity between trials, we defined a distance metric on the eye velocity at the initiation of the eye movement. We chose the sum of absolute differences (L_1_ norm) between the velocities in the 50 to 150 ms after the target motion onset. In the *k*^th^ shuffle (*k* = 1, 2…), we found the *k* most similar trials to each trial. We use the term “rank *k* correlation” to refer to the correlation calculated after allowing a shuffle of the *k* most similar trials. Since each trial is most similar to itself, the rank 1 correlation is simply the real, unshuffled correlation between the pair of neurons.

Figure 2A shows an example trial (black) alongside several trials from the same session that were similar to it. We shuffled the response of the trials in neuron 1, restricting the shuffle to the *k* most similar trials. Figures 2B and 2C illustrate an example of a *k*=3 shuffle. The reference trial in neuron 1 (black) could be shuffled either with the trial most similar to it (green) or the trial second-most similar to it (blue). In this example, it was randomly chosen to switch the reference trial with the second-most similar trial. Thus, in Figure 2C the blue trial replaces the black reference trial in neuron 1. We repeated this process for all the trials, resulting in a raster of shuffled trials. We then recalculated the correlation between the unshuffled neuron 2 and the shuffled data from neuron 1.

**Figure 2:**
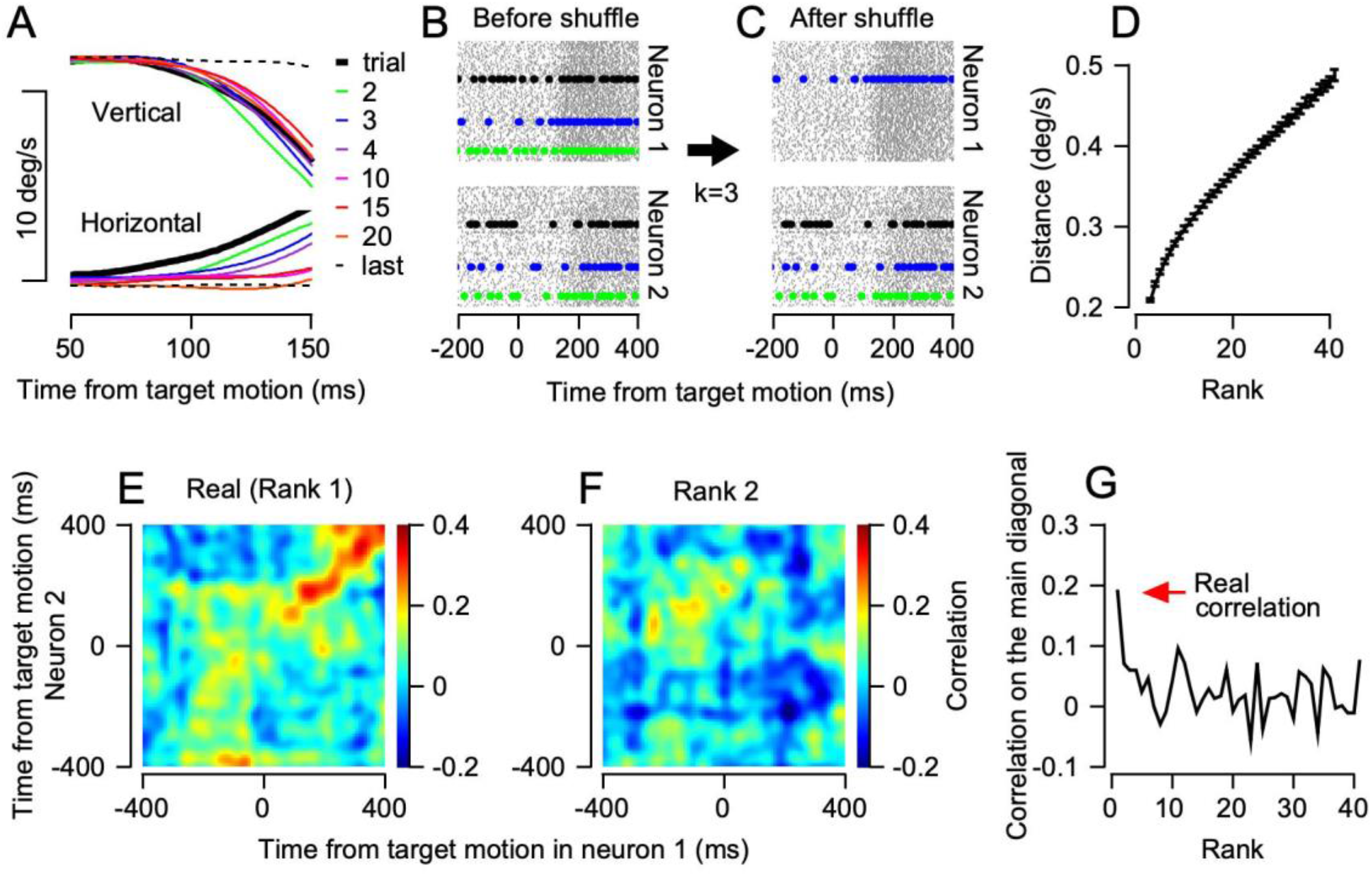
Matching similar behavior trials to estimate the behavior-related correlations. **A**, Eye velocity traces in the horizontal (bottom) and vertical (top) directions. The black trace represents a reference trial, and the color traces represent the *k*^th^-closest trial to it. The dashed line is the furthest trial in this session. **B**, Raster plot of an example pair of neurons (same as Figure 1D) before shuffling. The black line represents the reference trial, the green line represents the trial with the most similar behavior to the reference trial, and the blue line represents the second-most similar trial. **C**, The same neurons after the reference trial in neuron 1 was switched with the second-most similar trial. **D**, Distance in the behavior between shuffled trials as a function of rank, averaged over all sessions. **E**, The real (rank 1) correlation matrix of the example neurons. **E**, The correlation of the same pair as in **E**, after a rank 2 shuffle (most similar trial). **G**, Correlation in the example pair as a function of rank. The real correlation (rank 1) is marked by the red arrow.

As we increased the rank and allowed trials to be shuffled with less similar trials, the distance between a trial and the matched trial increased considerably (Figure 2D). Therefore, in lower ranks the correlations were calculated after shuffles between trials in which the behavior was substantially more similar. It is important to emphasize that for ranks higher than 1 we did not allow trials to be shuffled with themselves. Thus, any correlation that remained after shuffling is due to the similarity of the neurons’ firing rates on different trials.

Figures 2E and 2F show the real (rank 1) and rank 2 correlation matrices for the example neurons in Figure 1D. When calculating the correlation matrix without shuffling, we observed that this pair of neurons was strongly positively correlated. When calculating the rank 2 correlation matrix, in which each trial in neuron 1 was replaced with the trial most similar to it, the positive correlations along the diagonal were lost (Figure 2F). To compare the correlations across ranks, we focused our analysis on the time around motion onset (50 to 150 ms), since this is the time we used in our distance metric and the time at which behavior variability is known to be most strongly linked to neural activity (Joshua and Lisberger, 2014; Medina and Lisberger, 2007; Schoppik et al., 2008). We found that the correlations were strongly reduced for all ranks except for the first (Figure 2G). The gap between the real correlation and the correlations after shuffling suggests that the co-variability in these example neurons has only a slight influence on the trial-by-trial variability in behavior.

### Behavioral similarity accounts for only a small fraction of the trial-by-trial correlation between neurons

We expanded our analysis to the population. In line with previous reports (Joshua et al., 2009; Kohn and Smith, 2005; Lee et al., 1998), we found that pairs of neurons in our sample tended to be more correlated on a trial-by-trial basis when their average responses were similar (Figure 3A). For each pair of neurons, we calculated the correlation between the peristimulus time histograms (PSTHs) in both task conditions. These PSTH correlations were positively correlated with trial-by-trial correlations (Figure 3A). We used this result to focus our analysis unbiasedly on a subpopulation of pairs of neurons with high trial-by-trial correlations, thus increasing the likelihood of detecting behavior-related correlations.

**Figure 3:**
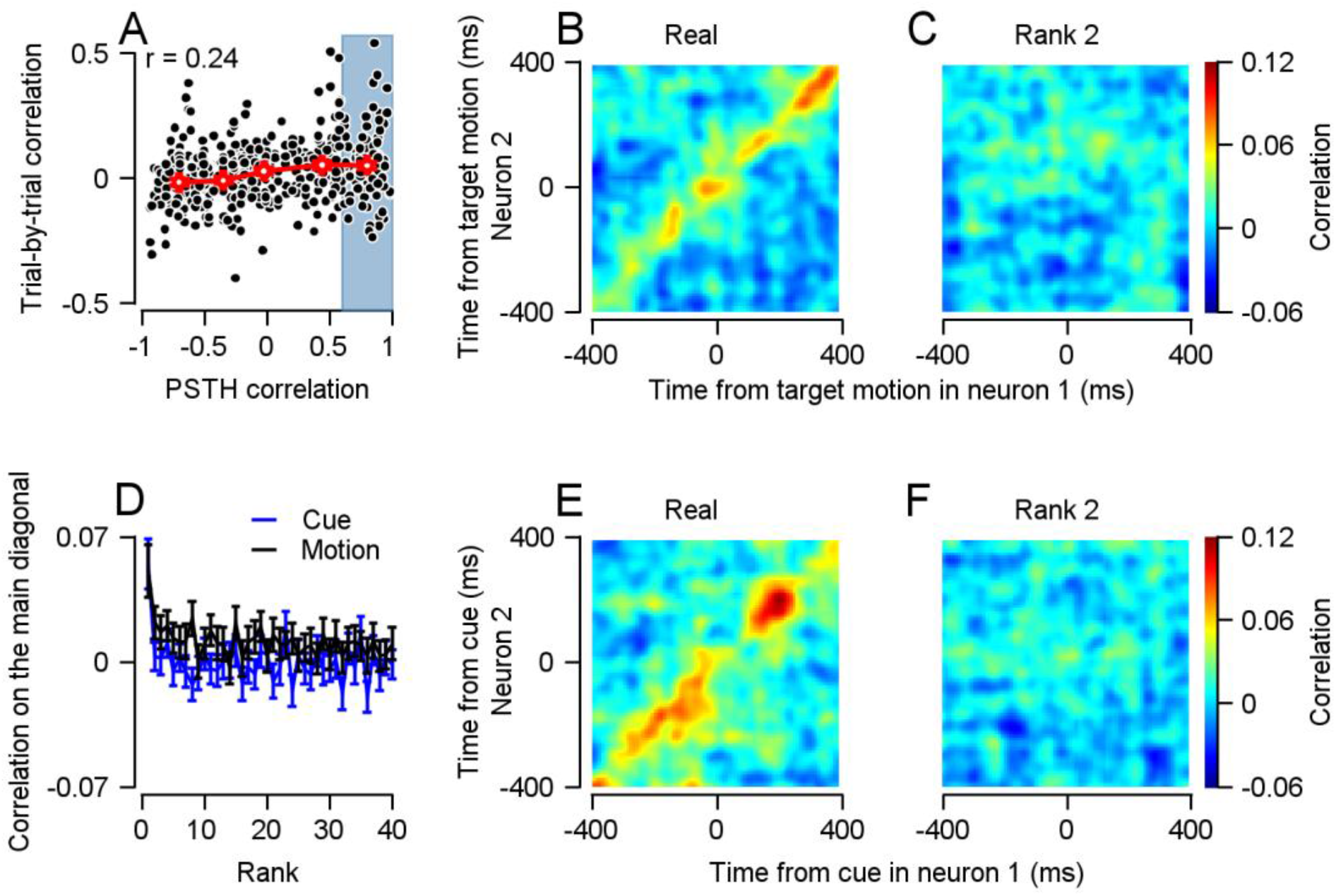
Estimating behavior-related correlations during motion onset and cue presentation in pairs with high PSTH correlations. **A**, The correlation between PSTH and trial-by-trial correlations (r=0.24, p=2.4*10^-7, n=470). Each dot represents a neuron pair’s PSTH correlation (horizontal) and trial-by-trial correlation (vertical). The red line represents the means of five quantiles of pairs, binned by PSTH correlation. Error bars in all the figures represent SEM. The blue rectangle marks the pairs that were used in **B** and **C. B** and **C**, Real (**B**) and rank 2 (**C**) correlation matrices for neurons within the top 20% of PSTH correlations (r>0.618, n=92) around motion onset. **D**, Average correlation in the 50 to 150 ms after target motion onset (black, signed-rank of rank 2: p=0.02, n=92) and the cue (blue, signed-rank of rank 2: p=0.5, n=92) as a function of rank. **E** and **F**, Real (**E**) and rank 2 (**F**) correlation matrices for neurons, within the top 20% of PSTH correlations (r>0.46, n=92) during cue.

The average trial-by-trial correlation matrix of pairs with the top 20% PSTH correlations (r>0.618) revealed a positive correlation around the main diagonal of the correlation matrix (Figure 3B). However, for the same pairs, this correlation declined sharply when calculating the rank 2 correlation matrix (Figure 3C). When we averaged the correlations along the diagonal 50 to 150 ms after motion onset, we found a significantly positive value of the rank 2 correlation matrix, although its magnitude was less than half of that of the real correlations (Figure 3D, black trace). As the rank increased, the correlations decreased even more. This result implies that in trials with similar behavior, the responses of pairs of neurons covary weakly. When repeating the analysis on the entire population, we found a similar decrease between the real and rank 2 average diagonal values, though the rank 2 correlations were not significantly positive (Figure S1).

The preparatory activity of neurons in the FEF has been linked to trial-by-trial differences in subsequent pursuit behavior (Raghavan and Joshua, 2017). We therefore extended our analysis to the cue period of the trials, focusing on pairs with high PSTH correlation during the cue. We calculated the different rank correlations around the cue presentation, using the pursuit behavior around motion onset in our distance metric as before. When aligning the correlations to the cue, we found positive correlations along the diagonal, both before and after the appearance of the colored targets (Figure 3E). We noticed a slight decrease in the correlation immediately after the appearance of the visual stimuli (Figure 3E, approximately 200 ms after cue onset) which is consistent with previous reports (Churchland et al., 2010; Kohn and Smith, 2005). The positive correlations did not appear in the rank 2 correlation matrix (Figure 3F). The average correlation along the diagonal 50 to 150 ms after the cue were close to 0 for all ranks larger than 1 (Figure 3D, blue trace). Thus, the correlations in the cue period were strongly attenuated even when the trials were shuffled by behavior similarity.

We verified that our results were not due to our choice of distance metric. We obtained similar results using the L_2_ norm (the sum of squared distances; Figure S2). FEF neurons code for eye movement acceleration as well as velocity (Mustari et al., 2009). We therefore included the eye movement acceleration in our analysis and verified that disparity between the real and rank 2 correlation matrices does not stem from the coding of acceleration (Figure S3). Lastly, we verified that the variable latencies of the corrective saccades after the initial pursuit do not explain the correlated activity (not shown).

The analysis displayed in Figure 3 focuses on subsets of positively correlated neurons (n=92). In this sample, we only found a small behavioral component in the trial-by-trial correlations. We used the fact that our analysis can be extended to neurons recorded at different times (see methods) to augment our sample. In these larger samples, we found clear shuffled correlations during movement (Figure S4), confirming that during movement part of the correlation is related to the behavior. However, they were small when compared to the real correlations that were obtained from simultaneously recorded neurons. The average shuffled correlations were comparable to what we observed in our smaller sample of simultaneously recorded neurons, suggesting that our samples were representative.

### Simulated data with shared population variability predicts a continuous decrease with rank

So far, we have relied on the notion that common coding of variable behavior should be reflected in the correlation between neurons. We next formalized this notion to verify that the shuffled correlations correctly estimated the behavior-related correlations. Our method does not assume a specific model of the FEF, we therefore simulated data using different plausible models. These models assume different relationships between neural activity and behavior. However, in each model, we can formally define behavior-related correlations. We used these models to study the factors that contribute to the pattern of correlation across ranks.

We began by simulating a simple one-dimensional network based on the model presented in Figure 1A. Although simple, this model captured the correlation between behavior and neural activity in the cerebellum and brainstem (Joshua and Lisberger, 2014; Medina and Lisberger, 2007). Formally, the firing rate *r*of the neurons in each trial was simulated as:

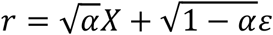

where, *X*∼*N*(0,1)was sampled once for each trial and shared across all neurons, *ε*∼*N*(0,1)was sampled for each neuron independently and *α*is a parameter between 0 and 1. In this network, the variance of each neuron is composed of a shared fraction (*X*, represented by the blue part of each neuron in Figure 1A) and an independent fraction (*ε*, red part). The behavior is the average activity of all neurons which equals to the shared component at the limit of a large number of neurons. In this simple network, any correlation between neurons contributes to the behavior. The correlation between neurons is equal to *α*.

We simulated the network for different values of *α*(between 0 to 0.05) and calculated the different rank correlations (Figure 4A). We found that the correlations for the simulated data decreased with rank, and we did not observe any gap between the real and rank 2 correlations. Rather, we found the decrease with rank to be almost linear, and as the ranks approached 1, the value of the correlations became close to *α*. This continuous decrease contrasts with the disparity between the shuffled and the real correlations in our data (Figure 3). This shows that, at least for this basic network, matching behavior similarity can approximate the fraction of correlations that contribute to behavior. However, this one-dimensional simulation is simplistic. To test whether these properties would still hold when we used a more complicated model, we applied our analysis on data generated from simulations of different plausible models of the FEF.

**Figure 4:**
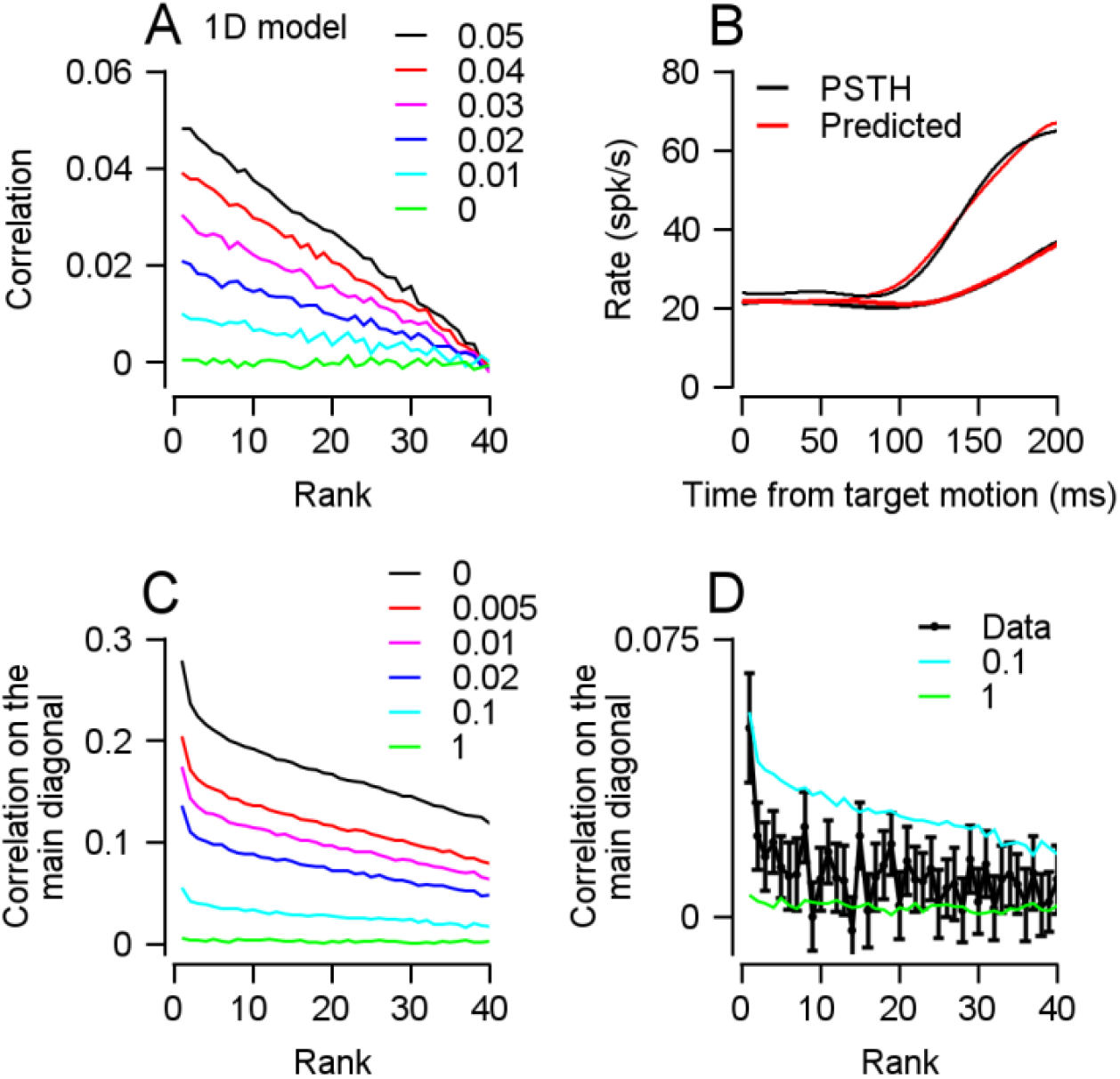
One-dimensional and kinematic model simulations show that behavior-related correlations can be estimated using the shuffling method. **A**, Correlation as a function of rank for simulated one-dimensional data. The different colors represent the values of *α*which determines the fraction of shared noise. **B**, Linear kinematic model fits for the two example neurons in Figure 1 (R^2^>0.98). **C**, Average correlation in the 50 to 150 ms after target motion onset as a function of rank for the kinematic linear model simulations (n=96). The different colors represent the degree of independent noise added to the simulation (see Methods). **D**, Zoom in on the same simulations as in **C**, along with the data from target motion onset (Figure 3D).

The relationship between rank and correlation may depend on the dimensionality of the behavior. The inclusion of different directions of movement (e.g., both horizontal and vertical) and the movement dynamics in time increase the dimensionality of the behavior. We therefore tested whether the shuffling method would correctly estimate behavior-related correlations when the behavior has the same statistics as the behavior in our task. We simulated the neural responses while using the actual behavior we recorded. We used a model that relates the eye movement kinematics to the neural response. In some of the FEF neurons, a linear relation appears to hold between eye kinematics and the firing rate (Mustari et al., 2009; Schoppik et al., 2008). To estimate this relation, we fitted a linear model between the average velocity and acceleration and the PSTHs of neurons. We focused on neurons with a high goodness of fit (R^2^>0.9, n=92; examples in Figure 4B). The model allowed us to calculate the predicted firing rate on each trial, based on the behavior alone. Thus, any correlation between these simulated neurons would be due to the similarity in the neurons’ responses to the same behavior. We added independent noise to the prediction of the model and applied our analysis.

Unlike in our neural data, the real and rank 2 correlations in the simulations were similar to one another, regardless of the amount of independent noise (Figure 4C). We only found a small disparity between the real and rank 2 correlation that did not resemble the large gaps we found in the actual data. Figure 4D compares the motion-aligned correlations from Figure 3D to the simulated correlations. The size of the disparity is much larger in the data than in the simulations that were matched to have the same rank 1 correlations. Thus, when considering higher-order data and the actual covariance structure of pursuit behavior in our task, we would expect to find substantial shuffled correlations. This implies that the disparity between the rank 2 and real correlation of the data is not a property of the behavior recorded in our task. Since some sessions had more than 40 trials, the rank 40 shuffle was not unconstrained in some neurons. Therefore, the correlations were slightly positive even in rank 40. However, when we averaged the correlations in the last rank that we could calculate for each neuron, we found the correlations reached 0 (not shown).

### Simulated population activity with plausible decoders predicts a continuous decrease with rank

The kinematic model simulation accurately represented the behavior in our data. However, it does not specify an explicit decoder of the population activity into behavior. To determine what factors could explain the gap between the real and the shuffled correlations in the data (Figure 3), we used a generative model of the population and subsequent behavior. We simulated a model of the FEF in which the neurons are directionally tuned and then read out by a population vector decoder. This model was suggested for cortical motor areas in similar tasks (Georgopoulos et al., 1986).

Figure 5A illustrates this model. Each neuron in our simulation has a preferred direction (PD) represented by its location on the ring. The correlation between neurons decreases with the distance of their PDs, specifically:

**Figure 5:**
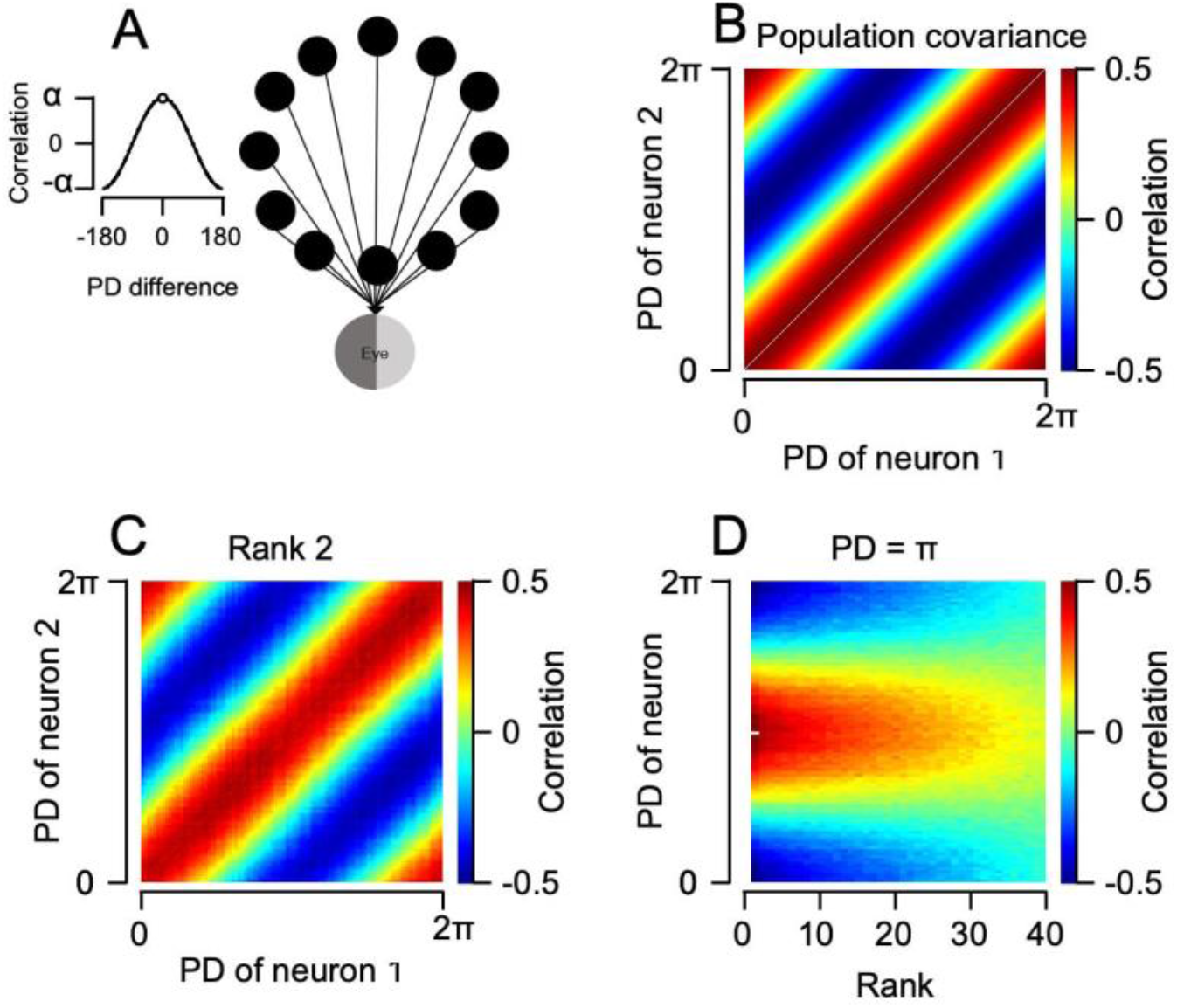
Population vector model simulations show that behavior-related correlations can be estimated using behavior similarity. **A**, Population model in which the covariance of neurons increases with the similarity in their PDs. **B** and **C**, The real (**B**) and rank 2 (**C**) correlation matrices between neurons with different PDs in the simulated population. **D**, For the neuron with a PD of *π*, the matrix illustrates the different rank correlations (horizontal) with neurons of different PDs (vertical). We omitted the real correlation of the neuron with itself (**B** and **D**) to highlight the other values in the matrix.

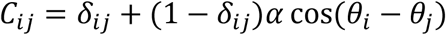

where *C*_*ij*_ is the correlation between neuron *i* and neuron *j, α*is a parameter that determines the maximal correlation between two different neurons *i*and *j, θ*_*i*_is the PD of neuron *i, θ*_*j*_ is the PD of neuron *j* and *δ*_*ij*_is the Dirac delta function (insert to Figure 5A). We set *α* =0.5since lower values had similar results but were more dominated by noise. Figure 5B shows the population correlation matrix *C* we used to simulate the population. Each point in the matrix represents the correlation *C*_*ij*_between two neurons whose PD is given by the horizontal and vertical positions. Note that the matrix in Figure 5B shows the correlation between all pairs of simulated neurons, and differs from previous correlation matrices (e.g., Figure 1E) which represent the average correlations of pairs of neurons at different times in the trial.

Each neuron contributes to a two-dimensional behavior (equivalent to horizontal and vertical movement). The behavior is calculated using the population vector decoder and is defined as:

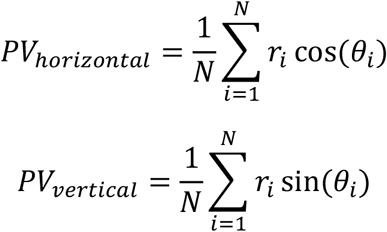

where *r*_*i*_is the rate of neuron *i* and *N* is the size of the population.

Although the relation between firing rate and behavior is complex, positively correlated neurons contribute similarly to behavior. This is guaranteed by the fact that positively correlated neurons have similar PDs so that their firing rates are weighted similarly by the decoder. In contrast, negatively correlated neurons have opposite PDs and thus contribute to behavior in opposite directions. We therefore expected that when shuffling behaviorally similar trials, the correlations (both negative and positive) would still be apparent.

As expected, the rank 2 correlation matrix was very similar to the real correlation matrix, both in its structure and in its values (Figure 5C). This is a marked departure from the actual data in which the correlations were strongly attenuated at ranks larger than one (Figure 3). The shuffled correlations depended on the distance between neurons PDs as demonstrated in Figure 5D. Figure 5D shows the correlations for a neuron with a PD of *π* with the rest of the population across ranks. The vertical axis represents the PDs of the other neurons, and the horizontal axis represents the rank. As the rank increases, the correlations approach 0 for both the positively and negatively correlated pairs. Thus, in this more complex model, the shuffling method can approximate the contribution of correlations to behavior.

### The interplay between the correlation matrix structure and the decoder determines the propagation of correlations to behavior

In the three simulations we presented so far, the correlations decreased with rank continuously and the real and rank 2 correlations had similar values. However, our analysis of FEF neurons shows a gap between the real and shuffled correlations (Figure 3). We used the population vector model to investigate possible mechanisms to explain this gap.

A possible explanation for the gap between the real and shuffled correlations is that parts of the neural correlations do not propagate to the behavior due to the interaction between the correlation structure and the decoder. In the cases described so far, the population was cosine correlated and the population vector decoder had a similar shape – a sinusoid with the same frequency. This is a unique case, in which the decoder assigns similar weights to highly correlated neurons. In the more general case of a population that is not cosine correlated, the mismatch between the decoder and the correlations might attenuate the correlations that propagate to the behavior. For example, for a uniformly positively correlated population, the population vector decoder will cancel out the correlations by subtracting the responses of neurons with opposite PDs. In more general terms, correlations will be averaged out when similar responses in neurons are weighted differently by the decoder.

We formalized this intuition mathematically and presented a detailed analysis in the Appendix. This formalization yielded interesting predictions about behavior-related correlations. Briefly, using Fourier transform, any population activity can be decomposed into a weighted sum of orthogonal vectors corresponding to different frequency sinusoids. The structure of the correlation determines these Fourier weights. The population vector decoder attenuates all frequencies resulting from the correlation structure except the one that corresponds to a sinusoid with the same frequency. Intuitively, the population vector filters out any frequency in the population that does not correspond to a sinusoid with a single cycle.

Thus, we can formally define behavior-related correlations as correlations that are not attenuated by the decoder. In the case of the population vector, the behavior-related correlations contain a single frequency. Consequently, even when the structure of the correlation is not cosine tuned, the behavior-related correlation will still be similar to the correlation matrix of the cosine tuned correlations (Figure 5C). Moreover, the magnitude of the behavior-related correlations depends on the weight of the single unattenuated component and can be calculated given the correlation structure.

The shuffling method correctly captured the analytically calculated behavior-related correlations. To demonstrate this, we varied the structure of the correlation, shifting it further and further away from the cosine correlations case (Figure 6A), and applied the shuffling method to data generated from the different populations. Figure 6B shows the rank 2 correlation matrix for a simulation that has a very different correlation structure from cosine correlations (cyan in Figure 6A). The rank 2 correlations were attenuated and did not reach the maximal real correlations in the population, as can been seen by the lack of bright red and blue in the correlation matrix. However, the structure resembled the one in Figure 5C, demonstrating that the qualitative shape reflects the one frequency that propagates through the population vector decoder. Figure 6C shows the correlations for a neuron with a PD of *π* with the rest of the population in the different ranks in the same simulation. The change in correlation structure had a similar effect as downstream noise. The shuffled correlation dropped sharply toward 0 and remained low at higher ranks. As the correlation structure departed further from a cosine, the rank 2 shuffled correlations became lower, though the real correlations remained high (Figure 6D, colors correspond to the correlation structures in 6A).

**Figure 6:**
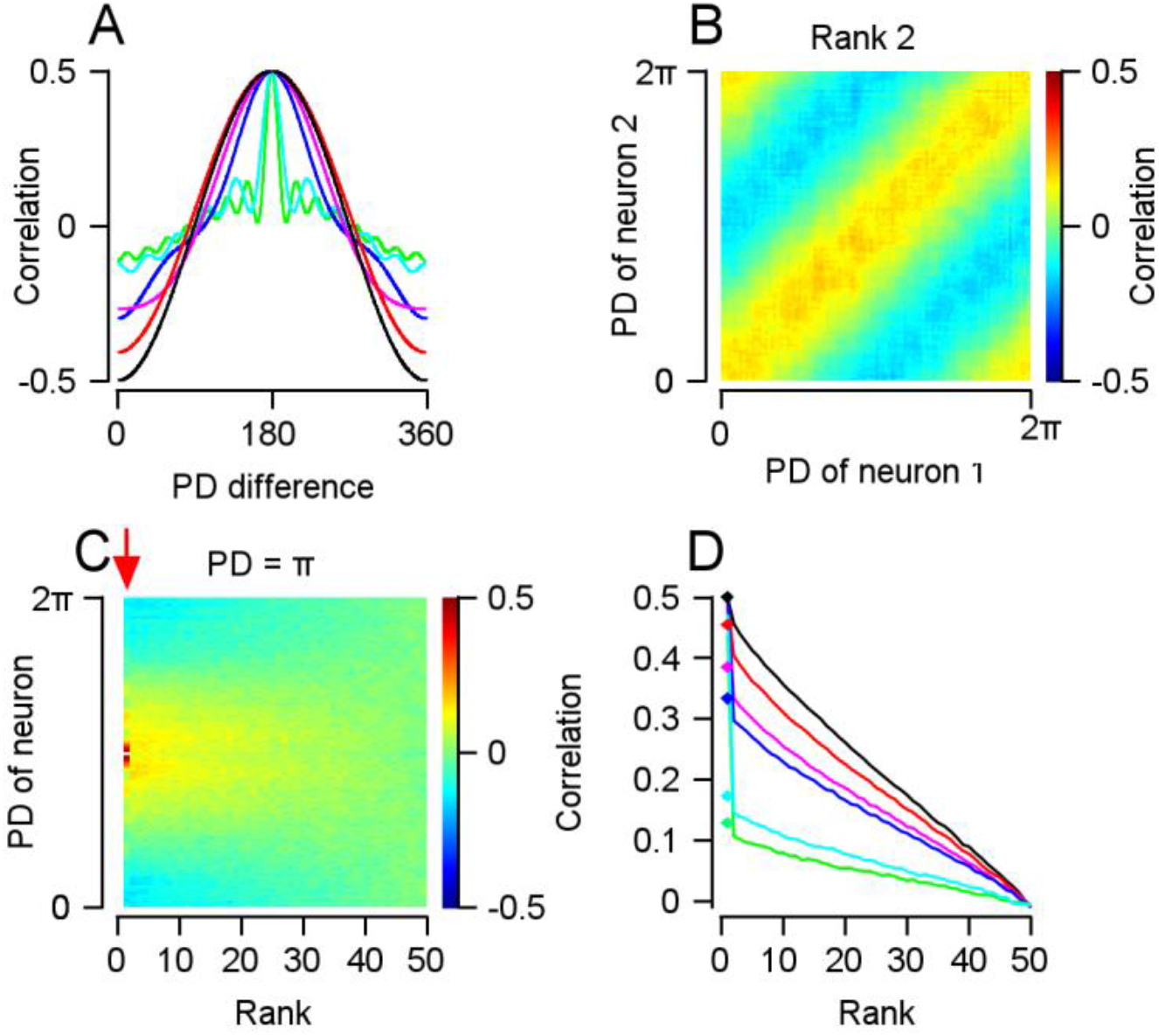
Interplay between the decoder and the correlation structure can cause a discrepancy between real and shuffled correlations. **A**, Different correlation structures were produced by adding an increasing number of frequencies (see Methods). **B**, The rank 2 correlation matrix produced in a simulation, using the correlation structure shown in cyan in **A. C**, For the neuron with a PD of *π*, the matrix illustrates the different rank correlations (horizontal) with neurons of different PDs (vertical). In this simulation we used the correlation structure shown in cyan in **A. D**, Correlation as a function of rank for simulated neurons with adjacent PDs for simulations with the corresponding correlation structures shown in **C**. We omitted the real correlation of the neuron with itself (**A** and **E**) to highlight the other values in the matrix.

We analytically calculated the value of the behavior-related correlations (diamonds in Figure 6D) and compared these values to the shuffled correlations. The shuffled correlations approached the value of the analytical behavior-related up to rank 2. When we increased the number of simulated trials per neuron, the rank 2 values reached analytical behavior-related correlations almost exactly (not shown). We repeated the same simulations for a lower value of *α* and obtained similar results (Figure S5). Thus, our method can correctly estimate behavior-related correlations in different population correlation structures.

Another possible explanation for the gap between real and rank 2 correlation is downstream noise. This noise can obscure the motor command from the FEF by increasing the behavioral variability independently of FEF activity (Medina and Lisberger, 2007). To investigate the influence of downstream noise on the correlations across ranks we simulated downstream noise as variability in the behavior that was independent of the output of the simulated FEF (see methods). The addition of downstream noise to the simulation resulted in a disparity between the real and shuffled correlations (Figure S6). The size of this disparity increased with the amount of downstream noise. This pattern of correlations across ranks is similar to that in our FEF data, suggesting that a large fraction of downstream noise is a possible explanation for the gap observed in the FEF data. Nevertheless, multiple studies have shown that most of the variability in the initiation of pursuit is caused by sensory components likely located upstream to the FEF (Hohl et al., 2013; Lee et al., 2016; Osborne et al., 2005). Furthermore, areas downstream to the FEF in the cerebellum, brainstem and motor periphery add only a small amount of noise at motion onset (Joshua and Lisberger, 2014; Medina and Lisberger, 2007). We therefore prefer other explanations that are consistent with both the current and previous findings.

Throughout these analyses, we showed that correlations that consist of frequencies different from the decoder of neural activity to behavior do not propagate to behavior. This provides a plausible explanation for the drop in correlation in the FEF data when shuffling behaviorally similar trials. Overall, these simulations demonstrate that when correlated FEF activity prorogates to behavior, we would expect a continuous decrease in correlations with rank.

## Discussion

We present a novel method to decompose correlations in the FEF into two factors: correlations that affect behavior and correlations that are independent of behavior. We found that only a small fraction of the correlations during movement and none of the cue correlations remain when shuffling the data, even when applying the strictest constraints on the similarity in pursuit behavior. This suggests that only a small fraction of the correlated noise in the FEF affects eye movements.

Several lines of evidence show that our method can detect behavior-related correlations. First, we found a clear correlation in behaviorally similar trials when we extended our analysis to include pairs of neurons not recorded at different times (Figure S4). In a different approach, we used several models of the FEF to simulate data and explore the different factors that can affect the correlation pattern. When all the correlations were expressed in behavior, we found that these correlations decrease gradually with rank (Figure 4A), even when the behavior had the same dimensionality as the behavior in our task (Figure 4C and D). Furthermore, our method correctly estimated behavior-related correlations in the presence of other correlations that do not affect the behavior (Figure 6). Thus, these simulations suggest that the behavior-related correlations can be estimated from the shuffled correlation. Importantly, our method makes modest assumptions about the FEF and does not require an explicit model of the relationship between activity and behavior. An unfitting model will result in a misestimation of the variance in firing rate that is related to behavior, and thus could bias the estimation of behavior-related correlation. The success of the simulations in capturing the behavior-related correlations demonstrates the generalizability of the approach as behavior-related correlations were estimated under different models of the FEF.

### Generalizing the shuffling methods to other behaviors and other recording areas

Our method is based on the increase of the distance in behavior with rank. As rank increases, trials become more separated in the behavioral space (Figure 2D). In low-dimensional space, this must be true. However, in high-dimensional behavioral space, it is possible to embed multiple trials with similar pairwise distances. In this case, all ranks larger than 1 will be practically equally distant. Thus, even if all the correlations are related to behavior, we would expect the correlation to increase only slightly as the rank approaches 2. Furthermore, as the distance of a trial from itself is always 0, there will be a gap between the distance of ranks higher than 2 and the distance in rank 1 that also is likely to be reflected in the correlations.

In the case of the pursuit behavior we studied here, the behavior is effectively low dimensional (Osborne et al., 2005), since vertical and horizontal behavior are correlated, and pursuit between adjacent time points is correlated as well. Thus, an apparently high-dimensional behavior is embedded in a low-dimensional space. This embedding is common to many behaviors (Sanger, 2000; Tresch and Jarc, 2009) and suggests that our method can be applied to many other domains. Practically, we note that compensating for higher dimensions requires a denser sampling of the behavioral space. Furthermore, the relation between the rank and distance (e.g., Figure 2D) can be used as a marker of the suitability of the shuffling method.

Our method can also be applied to other recording areas or other motor systems. As noted above, our method requires controlled and precise measures of behavior and a sufficient number of trials. This is often the case for recording in head-restrained animals so that it would be fairly straightforward to apply our method to recordings in other brain areas in the eye movement system, or other motor systems such as arm movement. Importantly, the number of pairs is not as critical as the control over behavior and the number of trials per neuron. We recorded 224 pairs simultaneously across 2 conditions. Since we had precise measures of behavior, this number was sufficient for reaching meaningful conclusions. A larger sample would of course improve the power of the tests, but we do not consider the number of pairs to be a limiting factor. Nevertheless, recording more neurons simultaneously could allow for the performance of additional tests, such as linking or dissociating higher-order patterns of activity and behavior.

### Quantifying the fraction of correlation that affects behavior

Our simulations illustrate cases in which we would expect either a continuous decrease in correlations with rank or a disparity between the real and shuffled correlations. The exact relationship between distance and correlation can be solved analytically for the one-dimensional case (the model in Figure 1A) but becomes complex for higher dimensions. However, our simulations show that for different models, the decrease with rank is approximately linear. Based on this finding, we extrapolated the correlation using a linear model of the decrease with rank to obtain a quantitative estimate of the fraction explained by behavior. For the pairs used in Figure 3, the extrapolated behavior-related correlations were 0.01 during movement (in comparison to the average real correlations of 0.051) and - 0.001 during the cue (in comparison to the average real correlations of 0.054). Thus, the behavior-related correlations comprised approximately 23% of the correlations during movement and do not explain the positive correlations during the cue.

The small extrapolated correlations suggest that the majority of the correlations were not pursuit-related. The residual part of the correlation could be attributed to the common representation of other internal or external variables. The FEF has been shown to be modulated by attention and by expected reward (Glaser et al., 2016; Lixenberg and Joshua, 2018). Another possibility is that the FEF does not only represent eye movement behavior but also additional motor behaviors. Recent work has shown a widespread and overlapping representation of different movements across the brain, which is inconsistent with the division of the brain into specialized modules controlling a single end-effector (Musall et al., 2019). In this case, defining the distance metric over the full behavior could show that the correlations are explained completely by the behavior. Regardless of whether the correlation is a result of other behaviors or internal processes, our analysis indicates that in the transformation to eye movements the correlations are attenuated.

### Conditions under which correlations fully propagate to behavior

The mathematical formulation of our population activity model indicates that the correlations fully propagate to the behavior only when the structure of the correlation is very simple and perfectly matches the decoder structure (Figure 6C-F). We reached this conclusion by assuming a population vector decoder, but similar conclusions can be drawn with more general assumptions. With any linear decoder, the behavior-related correlations can be calculated based on the overlap (or angle) between the decoder and the Fourier components of the correlations. The correlation will only be completely behavior-related in the singular cases where the decoders match the correlation structure perfectly. Furthermore, in the case of other possible non-linear decoders such as the maximum likelihood estimator of the eye movement speed and angle, the conditions under which correlations fully propagate to behavior are unique (not shown). Thus, even under more general conditions than the ones we simulated we would not expect all the correlations to propagate to behavior. Our simulation suggests that the decoder can act as a filter, allowing only specific patterns of activity to pass onward to behavior. This idea is in line with recent findings regarding the propagation of correlations in the visual system (Semedo et al., 2019).

## Methods

### Experimental Design

In this paper, we reanalyzed data reported in a published study (Raghavan and Joshua, 2017). The data were collected from two male rhesus macaque monkeys (*Macaca mulatta*) that had been trained to perform behavioral tasks and prepared for neural recording using techniques described in detail previously. All procedures were in strict compliance with the National Institutes of Health Guide for the Care and Use of Laboratory Animals and were approved in advance by the Institutional Animal Care and Use Committee at Duke University, where the experiments were performed. In brief, we implanted head holders to restrain each monkey’s head, and a coil of wire on one eye to measure eye position using the magnetic search coil technique (Judge et al., 1980). In a later surgery, we placed a recording cylinder stereotaxically over the frontal eye fields. We lowered up to five quartz-insulated tungsten electrodes into the caudal parts of the FEF to record spikes. For the detailed data analysis, we sorted spikes offline (Plexon Inc., USA).

When lowering the electrodes, we looked for neurons that responded during pursuit eye movements. After we characterized pursuit tuning, we subjected the monkeys to the main experiment protocol in which we introduced interleaved target-selection and single-target trials while manipulating the reward size. The characterization of the trial-by-trial fluctuations in neural activity and behavior required many repetitions of the same experimental condition. As the number of single-target trials was small we focused here on the two-target trials.

Each of the target-selection trials started with a bright white circular target that appeared in the center of the screen. After 500 ms of presentation, in which the monkey was required to acquire fixation, two peripheral colored targets appeared at 3° eccentric to the fixation target, along orthogonal directions. The monkeys were required to continue to fixate the white target in the center of the screen. The color of the target indicated the size of the reward that the monkey would receive if it tracked the target. After a random delay of 800-1,200 ms, the colored targets stepped to a location 4-5° eccentric to the center of the screen and then started to move toward the center of the screen at 30°/s, prompting the monkey to initiate a smooth pursuit eye movement. In different sessions, the targets moved along different axes but were always orthogonal to one another. Typically, the monkeys initiated a pursuit eye movement that was biased to the larger reward target, followed by a saccade that brought the eye position toward one of the targets. Online, we detected this saccade and blanked the target that was further from the eye after the saccade. At the end of the trial, the target stopped, and if the eye position was within a 2°x2° window around the target, the monkey received a juice reward.

### Data analysis

All analyses were performed using MATLAB (MathWorks). For the analysis, we included neuron pairs that were recorded simultaneously for more than 40 trials in a condition. We used only pairs recorded on different electrodes to avoid sorting errors that result from overlapping extracellular waveforms. We excluded trials with saccades in the 15 ms before target motion or in the 235 ms after target motion onset. After applying these criteria, we were left with 224 pairs of neurons, consisting of 188 single neurons. Eleven pairs were recorded in more than one session, we included these additional sessions in our analysis. Removing them did not alter our results.

To study the average properties of the neuron’s response in time, we calculated the PSTH at a resolution of 1 ms. We then smoothed the PSTH with a 15 ms standard deviation Gaussian window, removing at least 75 ms before and after the displayed time interval to avoid edge effects. Note that this procedure is practically the same as measuring the spike count per trial in larger time bins. We defined the PSTH correlation between two neurons as the Pearson correlation between the neurons’ PSTHs in the 220 ms after target motion onset or cue presentation concatenated over experimental conditions.

To calculate the (rank 1) trial-by-trial correlation matrix of pairs, we first calculated single-trial PSTHs. For efficiency reasons, we binned the PSTHs to 50 ms windows. We calculated the Pearson correlation over trials of the neurons’ PSTHs at different time points. When plotting the average correlation, we refer to the average of the main diagonal of the correlation matrix in the 50 to 150 ms after the target motion. As a control, we shifted the trials in one of the neurons one trial forward and recalculated the correlations in the same way (Figure 1F, gray histogram).

We smoothed the pursuit single-trial velocity traces with a 15 ms standard deviation Gaussian window, again removing at least 75 ms before and after the time interval we used for analysis to avoid edge effects. Changing the standard deviation of the Gaussian window did not cause consistent changes in the results. We calculated the distance between every two pairs of trials by concatenating the horizontal and vertical traces in the 50 to 150 ms after the target motion onset of each trial. This resulted in two vectors in a 202-dimensional space. When we included the acceleration in the distance metrics (Figure S3), these vectors were in a 404-dimensional space. We defined the distance as the L_1_ norm of the difference between these vectors.

When applying the shuffling method to a pair of neurons we randomly chose which neuron was subjected to shuffles (neuron 1), whereas the other neuron was not shuffled (neuron 2). When calculating the rank *k* correlations, we iterated over the trials recorded in neuron 1. For each trial, we sorted all other trials by their behavioral difference from that trial. We randomly chose a trial that was within the *k* most similar trials to it (but not the trial itself). We replaced the original trial with the behaviorally similar one. After iterating over all trials, we had a shuffled version of the neuron’s response. We used the shuffled response and the unshuffled response of the other neuron and proceeded as described above. We also used an exact shuffle, in which we calculated the rank *k* shuffle by shuffling with the *k*^th^-most similar trial without randomization. Using the exact shuffle, we reached the same conclusions as with the randomized shuffle. However, the exact shuffle is inferior to randomized shuffle since in large ranks outliers are matched to many trials, leading to a substantial amount of noise in the estimates of the correlations. When plotting the distance as a function of rank (Figure 2D), we plotted the average of the absolute differences. To linearly extrapolate the real correlations from the shuffled correlations we fit a robust linear regression model from distances to the shuffled correlations.

The shuffling method can be extended to neurons recorded at different times. To calculate rank 2 and higher correlations we do not need the activity of the neurons in the same trial, only similar trials. Therefore, the shuffled correlations can be calculated as long as the behavior in the session is similar. When neurons are not recorded simultaneously, we cannot compare shuffled correlations to the real correlations, but we can get an estimate of the behavior-related correlation sizes. When calculating correlations of rank 2 and higher for neurons that were not recorded simultaneously, we paired neurons that were recorded in sessions in which the targets moved along the same axis, as anisotropies could lead to large differences in the behavior in different directions.

### Simulations

#### One-dimensional model

We simulated 50,000 pairs of neurons in 40 trials each. In each trial, the behavior was simulated as a normal random variable *X* with mean 0 and variance 1. The neurons’ responses were simulated as 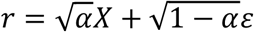 where *ε* was sampled from a normal distribution with mean 0 and variance 1 for each neuron independently and *α*was set to different values between 0 and 0.05. The distance between trials was defined as the L_1_ distance between the *X* s in the trials.

#### Linear kinematic model

To fit the model, for the two conditions in each neuron we first smoothed the PSTHs, the eye velocity, and the eye acceleration in the 220 ms after movement onset. We concatenated the two conditions and fit a linear regression model from the average behavior to the PSTH (four regressors: horizontal velocity, vertical velocity, horizontal acceleration, vertical acceleration). We estimated the model fit using R^2^. For pairs in which both neurons had a good fit (R^2^>0.9, 48 pairs out of 224), we used the coefficients obtained from the average response to predict the firing rate in single trials based on the behavior (Joshua and Lisberger, 2014; Lixenberg et al., 2020; Medina and Lisberger, 2007). We added independent normal random noise to the simulated single-trial firing rate that was proportional to the difference in trial-by-trial variance between the neurons and the model predictions (different colors in Figure 4C and D). We then performed the same analysis on these simulated single-trial firing rates as we did on real data. We repeated this 100 times for each of the pairs that had a good model fit and averaged them, to reduce the noise.

#### Population vector model

We repeated 100 simulations of a network of 1,000 neurons for 50 trials. The figures we show are the averages of these simulations. Each neuron was assigned a PD between 0 and 2*π* with equal intervals between two adjacent neurons. The simulated correlation of the neurons depended on their PD difference, as described in the Results section with α = 0.5. To simulate pseudo-random numbers with a specified correlation matrix, we used the Cholesky factorization on pseudo-random normally distributed neurons. The behavior in each trial was specified in the Results section, and the distance between the behavior in two trials was defined as the L_1_ norm of the difference between the 2-dimensional behavior vectors. When we set α = 0.05 (Figure S5) we increased the number of repetitions in the simulations to 1,000 to reduce the noise.

We added simulated downstream noise by adding noise to the behavior, independent of FEF activity. Specifically:

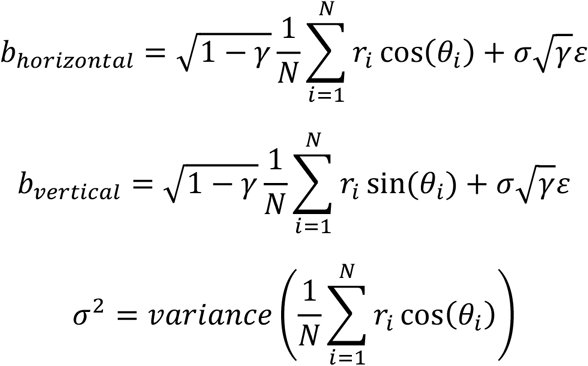

where *ε* is sampled from a normal distribution independently for the behavior and 0 ≤ *γ* ≤ 1 is a parameter that determines the fraction of behavior variance explained by FEF activity.

We varied the correlation function of the population by changing its frequency components. We added a constant to an increasing number of frequencies (in addition to the first) from low to high:

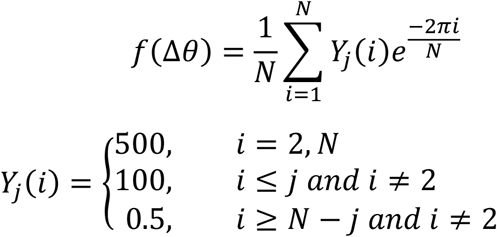

We then normalized the resulting function so that the maximal correlation was *α* and set the correlation of a neuron with itself to 1. The resulting correlation functions as shown in Figure 6C correspond to *j* = 0, 1, 3, 4, 11, 16 corresponding to black, red, pink, blue, cyan, and green.

## Appendix

The population activity vector *r* can be decomposed using the real Fourier transform:

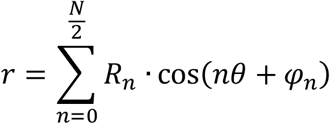

where *R*_*n*_ and *φ*_*n*_ are the amplitude and phase, respectively, of the real Fourier decomposition and *θ* is a vector containing the PDs of the neurons. In this notation, the normalization factors are contained in the amplitude term. The population vector decoder (cos(*θ*) and sin(*θ*)) is orthogonal to all Fourier components except the second, therefore

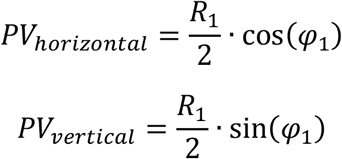

Thus, the population vector outcome in each trial is determined solely by the size of *R*_1_ and the phase shift in this component (*φ*_1_).

The covariance between two neurons is:

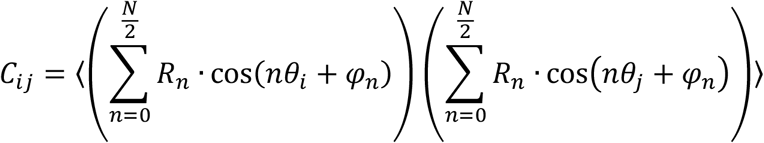

By using the orthogonality of the different components again, this is equal to:

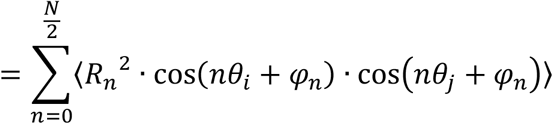

Using trigonometric identities, we can rearrange the equation:

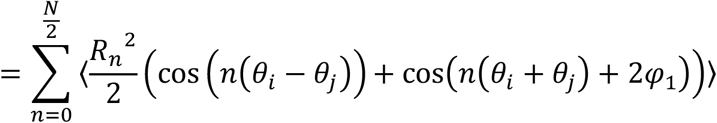

Since the distribution is multivariate Gaussian, the phase is uniformly distributed, and the expectation of its cosine is 0. In addition, since the correlation matrix is circulant, the phase and amplitude are independent. Therefore, the component containing the phase is averaged out:

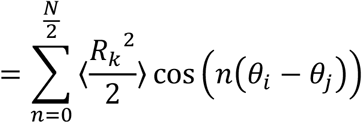

In the case of the population vector, we can use this decomposition to separate the covariance matrix into frequencies that propagate through the decoder from correlations that do not:

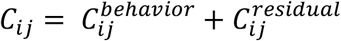

Where

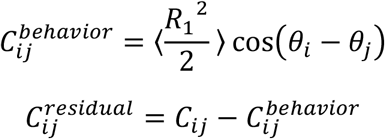

and *C*^*behavior*^is the behavior-related correlation matrix.

## Acknowledgments

We thank Lisberger S.G. for his support in the early stages of this project. We thank Burak Y., Lottem E., Rokni D., Shamir M., Harpaz M and Darlington T. for commenting on early versions of the manuscript. This work was supported by the European Research Council (755745) and the Israeli Scientific Foundation (380/17).

---

**Figure S1:**
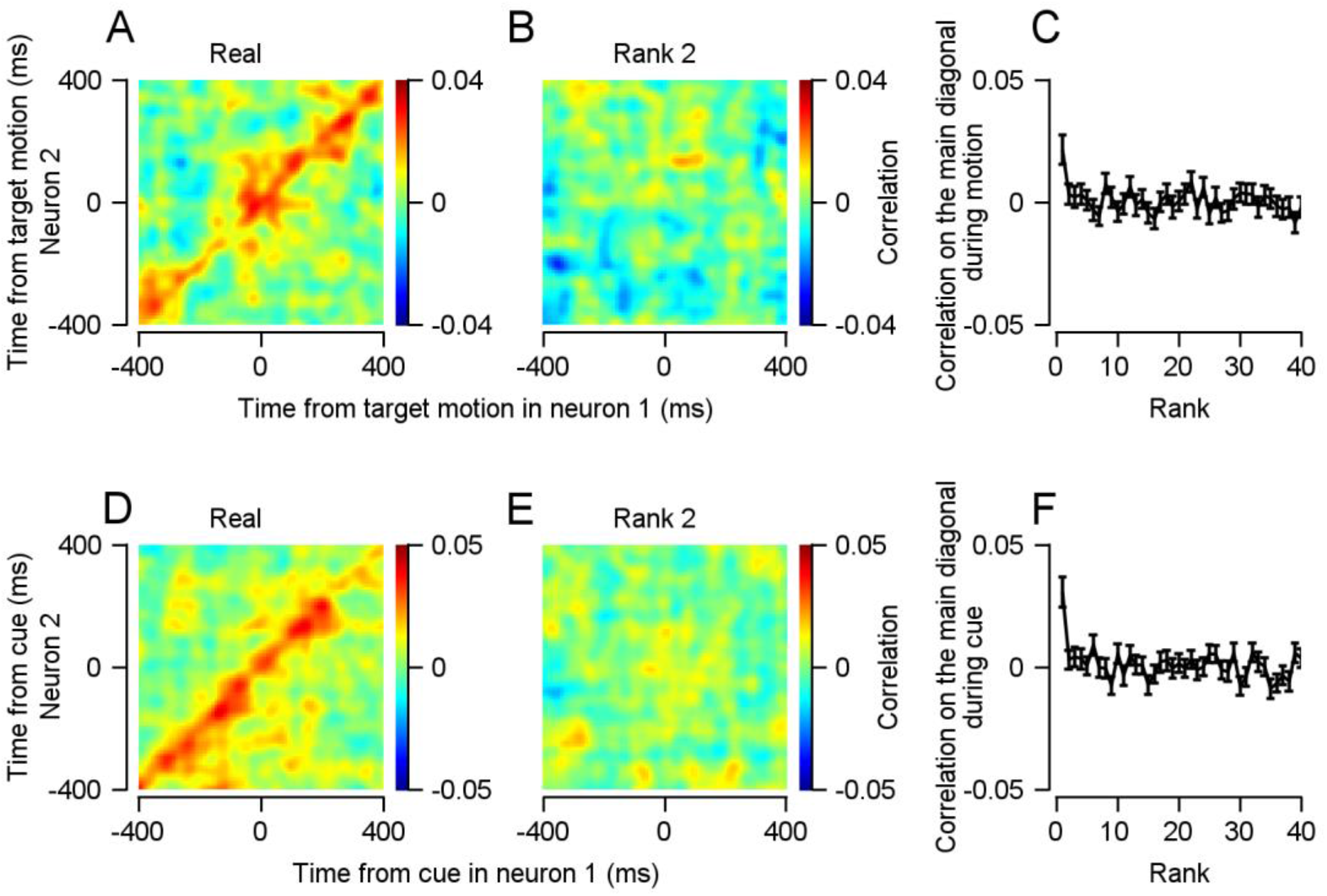
Estimating behavior-related correlations in the entire population yields similar shuffled correlations to the subpopulation with high PSTH correlations. **A, B**. Real (**A**) and rank 2 (**B**) correlation matrices for the entire population around motion onset. **C**, Average correlation in the 50 to 150 ms after target motion onset as a function of rank (signed-rank of rank 2: p=0.45, n=470). **D, E**. Real (**D**) and rank 2 (**E**) correlation matrices for the entire population during cue. **F**, Average correlation in the 50 to 150 ms after cue onset as a function of rank (signed-rank of rank 2: p=0.5, n=470).

**Figure S2:**
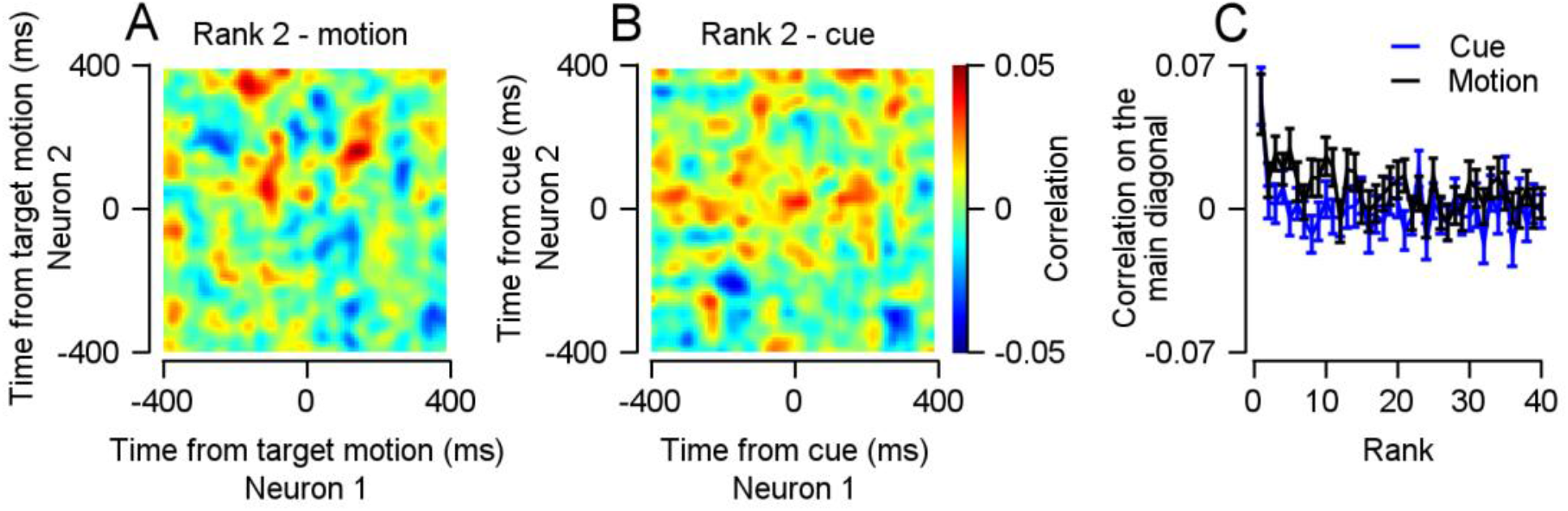
Measuring behavioral distance using the L2 norm yields similar shuffled correlations. **A** and **B**, The rank 2 correlation matrices of the same neurons as in Figure 3B (**A**) and in Figure 3E (**B**), using the L2 norm as the distance metric instead of the L1 norm. **C**, Average correlation in the 50 to 150 ms after target motion onset (black, signed-rank of rank 2: p=0.24, n=92) and after the cue onset (blue, signed-rank of rank 2: p=0.89, n=92) as a function of rank.

**Figure S3:**
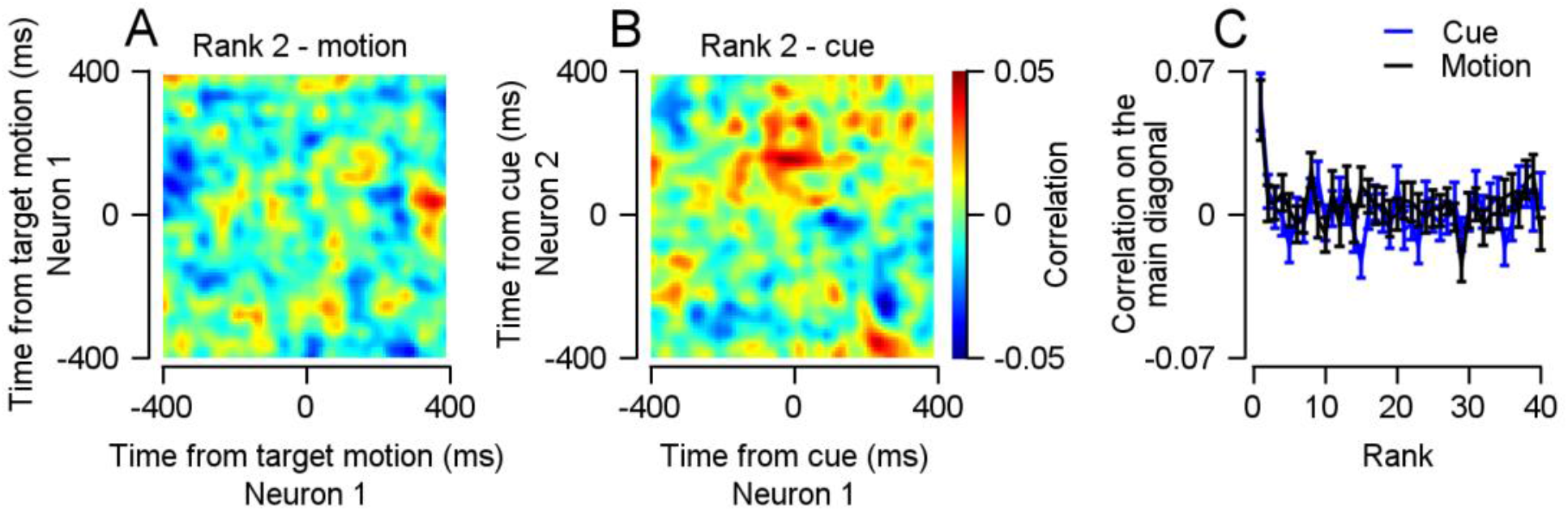
Including pursuit acceleration in the behavioral distance metric marginally increases the shuffled correlations. **A** and **B**, The rank 2 correlation matrices of the same neurons as in Figure 3A (**A**) and in Figure 3D (**B**), including acceleration in the distance metric. **C**, Average correlation in the 50 to 150 ms after target motion (blue, signed-rank of rank 2: p=0.68, n=92) and after the cue (blue, signed-rank of rank 2: p=0.3, n=92) as a function of rank.

**Figure S4:**
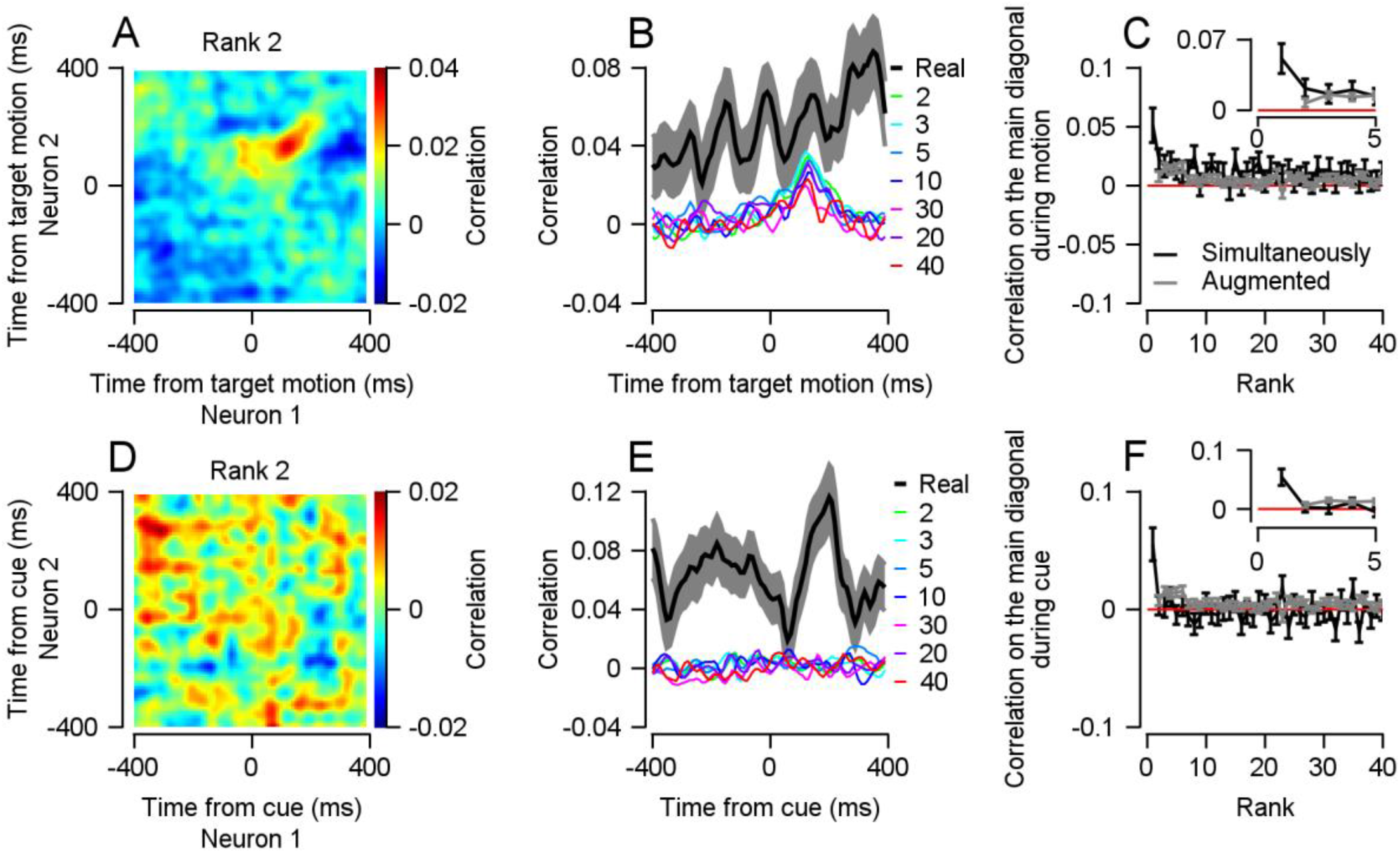
Estimating behavior-related correlations around motion onset in neurons recorded simultaneously or in different sessions. **A**, The rank 2 correlation matrix of neurons with matching PSTH correlations (r>0.618) to the neurons analyzed in Figure 3B, aligned to target motion onset. Note that the neurons in this figure were not necessarily recorded in the same session. **B**, The thin color traces represent the main diagonals of the different correlation matrices, with each color corresponding to one correlation matrix’s rank. The thick black line represents the main diagonal of the real correlation matrix in Figure 3B. **C**, Average correlation in the 50 to 150 ms after target motion onset as a function of rank for neurons that were not necessarily recorded together (grey; signed-rank for rank 2: p=1.3*10^-9, n=490) and for neurons that were recorded together (black, the same as in Figure 3D; signed-rank of rank 2: p=0.02, n=92). The thin red line represents 0 correlation. The insert shows ranks 1 to 5. The correlations in pairs recorded simultaneously were similar in size to the augmented sample. **D**-**F**, The same analysis as in **A**-**C** (grey; signed-rank for rank 2: p=0.65, n=360) for the neurons in Figure 3E aligned to cue onset (black, the same as in Figure 3D; signed-rank of rank 2: p=0.5, n=92).

**Figure S5:**
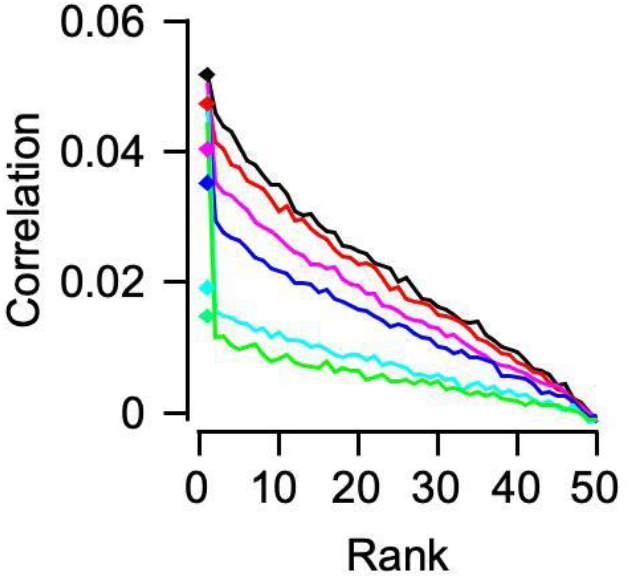
Estimation of behavior-related correlations in the population vector simulation is qualitatively similar when *α*is low. Same as Figure 6D, with *α*= 0.05 instead of *α*= 0.5.

**Figure S6:**
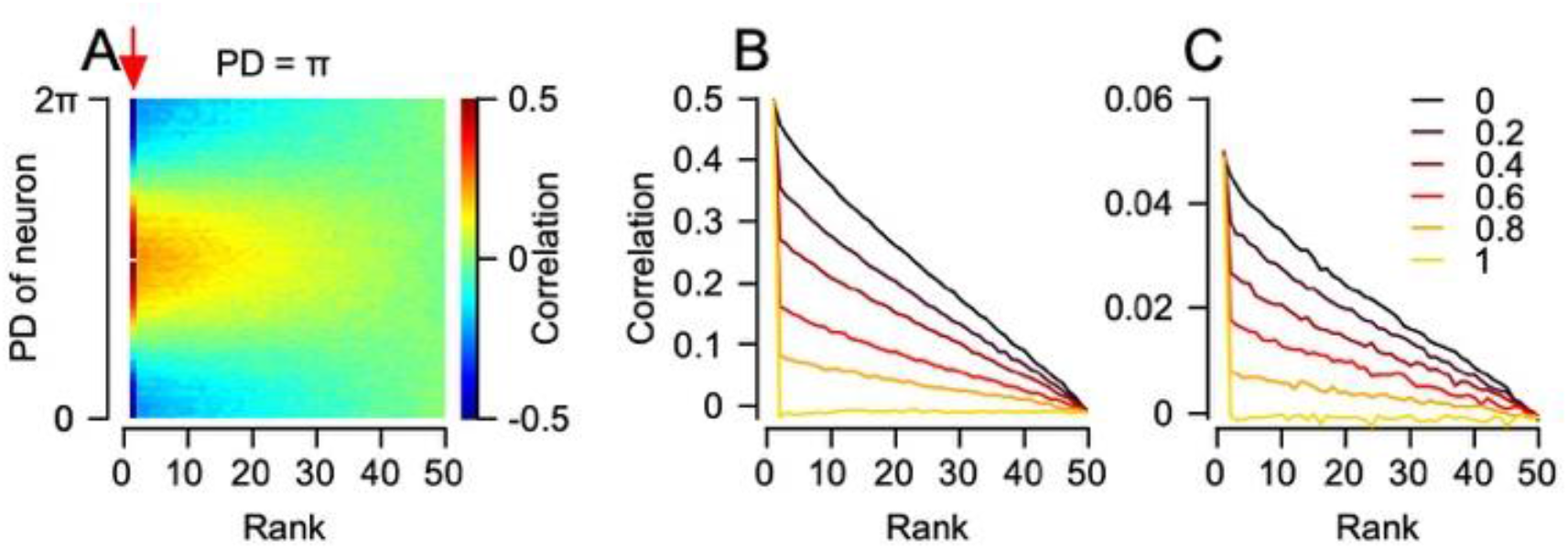
Downstream noise can cause a discrepancy between real and shuffled correlations. **A**, For the neuron with a PD of *π*, the matrix illustrates the different rank correlation (horizontal) with neurons of different PDs (vertical) when the fraction of downstream noise was 0.5 and *α*= 0.5. **B** and **C**, Correlation as a function of rank for simulated neurons with adjacent PDs. The different colors represent simulations with different fractions of downstream noise. *α*=0.5and **B** and 0.05 in **C**.

